# Characterization of osteoarthritis phenotypes by global metabolomic profiling of human synovial fluid

**DOI:** 10.1101/395020

**Authors:** Alyssa K. Carlson, Rachel A. Rawle, Cameron W. Wallace, Ellen G. Brooks, Erik Adams, Mark C. Greenwood, Merissa Olmer, Martin K. Lotz, Brian Bothner, Ronald K. June

## Abstract

**Objective:** Osteoarthritis (OA) is a multifactorial disease with etiological heterogeneity. The objective of this study was to classify OA subgroups by generating metabolic phenotypes of OA from human synovial fluid.

**Design:** *Post mortem* synovial fluids (n=75) were analyzed by high performance-liquid chromatography mass spectrometry (HPLC-MS) to measure changes in the global metabolome. Comparisons of healthy (grade 0), early OA (grades I-II), and late OA (grades III-IV) donor populations were considered to reveal phenotypes throughout disease progression.

**Results:** Global metabolomic profiles in synovial fluid were distinct between healthy, early OA, and late OA donors. Pathways differentially activated among these groups included structural deterioration, glycerophospholipid metabolism, inflammation, central energy metabolism, oxidative stress, and vitamin metabolism. Within disease states (early and late OA), subgroups of donors revealed distinct phenotypes. Phenotypes of OA exhibited increased inflammation (early and late OA), oxidative stress (late OA), or structural deterioration (early and late OA) in the synovial fluid.

**Conclusion:** These results revealed distinct metabolic phenotypes of OA in human synovial fluid, provide insight into pathogenesis, represent novel biomarkers and assist in developing personalized interventions for subgroups of OA patients.

## Introduction

Osteoarthritis (OA) affects over 250 million individuals worldwide and is associated with an annual economic burden of at least $89.1 billion [1]. OA is the most common joint disease characterized by pain and loss of function resulting from the breakdown of the articular cartilage [2]. Pathologically, OA joints exhibit cartilage damage, osteophyte formation, subchondral bone sclerosis, and varying degrees of synovitis [3]. Altered joint metabolism, inflammation, increased joint loading, joint injury, and other factors contribute to the development of OA [4-8].

This multifactorial nature of OA contributes to a broad variation in presentation of symptoms, progression of disease, and response to treatments. In addition to the multiple contributing factors, the trajectory of OA prognosis is highly variable. Some patients rapidly progress into severe stages of disease, whereas others remain relatively stable for decades [9-12]. Similarly, the perception of pain is also variable, with some patients experiencing minimal pain despite obvious joint space narrowing and others experiencing extreme pain with minimal joint space narrowing. OA was recently described as having multiple phenotypes in which subsets of disease characteristics drive differences between subgroups of patients with distinct OA outcomes [8]. However, more data are needed to define these phenotypes.

OA heterogeneity poses many challenges for understanding pathogenesis, facilitating diagnosis and therapeutic interventions [13-15]. Defining phenotypes of OA is important for many reasons. First, this would provide insight into factors that contribute to the development of these distinct phenotypes [8]. Secondly, it would allow for development of targeted treatments for specific subgroups of OA [8]. Finally, given the heterogeneity of OA, defining phenotypes is crucial for identifying biomarkers for early diagnosis across all phenotypes or within specific subgroups once identified.

Metabolomics is a promising method for distinguishing phenotypes. Metabolomics analyzes large numbers of small-molecule intermediates [16]. Changes in the metabolome occur rapidly and reflect the overall biological response from changes in the genome, transcriptome, and proteome [17]. Metabolomic profiling generates a phenotype that characterizes functional cellular biochemistry [16, 17]. Global metabolomics is promising because it produces a global view of the metabolome with minimal bias. By focusing on all metabolite features in the sample, this analysis develops a network of pathways that illustrate metabolic perturbations with disease. Therefore, global metabolomic profiling is not only beneficial for identifying specific metabolites as potential biomarkers as demonstrated previously [18], but also providing insight into the underlying mechanism of disease.

The SF is an ultrafiltrate of the plasma containing additional molecules produced by the cells in joint tissue. SF provides lubrication between the articular cartilage surfaces and eliminates metabolic waste. The SF is in direct contact with other OA-affected tissues (*i.e.* articular cartilage, synovium, etc.) and will reflect local changes with disease [19]. This makes the SF a promising biofluid for phenotype identification given the heterogenous pathology of OA in the joint.

The objective of this study is to apply our established LC-MS-based global metabolomic profiling method to generate metabolic phenotypes of SF from donors across all stages of OA (grades 0-IV). By characterizing global metabolomic profiles of early and late OA, this study seeks to (1) identify differences in metabolic pathways throughout disease progression from healthy to late stage disease, and (2) classify patients within early and late OA into subgroups representative of potential OA phenotypes. To our knowledge, this is the first study to perform global metabolomic profiling of SF from donors with early and late stage OA to investigate metabolic perturbations throughout disease progression.

## Methods

### Human Synovial Fluid

*Post mortem* SF samples (n=75) from knee joints were used for this study under an IRB exemption. Joints were graded based on severity of changes in the knee cartilage surfaces using the Outerbridge scoring system which grades joints from 0-IV based on macroscopic cartilage pathology[20]. The distribution of OA knees was as follows: grade 0 (n=7), grade I (n=28), grade II (n=27), grade III (n=13), and grade IV (n=4). SF samples were grouped in three cohorts: healthy controls (grade 0; n=7), early OA (grades I-II; n=55), and late OA (grades III-IV; n=17). These samples include both sexes and a variety of ages (Table 4.1). SF was frozen at −80°C until analysis. All samples were de-identified and blinded prior to mass spectrometry and data analysis.

### Donor Demographic Information

Age, sex, and OA grade were included for all donors (Table 1). Additional clinical data available for some but not all donors included donor height and weight, cause of death, pre-existing medical conditions, and history of OA.

**Table 1.** Descriptive statistics for donor population. Descriptive statistics of donor population for each cohort including age, sex (as male % population), and BMI. All means are reported as mean +/-standard error. BMI was unavailable for some donors (BMI=body mass index).

| Cohorts | Healthy | Early OA | Phenotype E1 | Phenotype E2 | Late OA | Phenotype L1 | Phenotype L2 |
| --- | --- | --- | --- | --- | --- | --- | --- |
| Age | 35 ± 4.796 | 55.47 ± 2.163 | 56.45 ± 2.781 | 54 ± 3.496 | 68.53 ± 3.868 | 66 ± 4.637 | 78.88 ± 4.006 |
| Sex (% male) | 57.14% | 52.73% | 54.55% | 50% | 41.18% | 27.27% | 66.67% |
| BMI | 24.2 ± 2.524 | 27.82 ± 0.9789 | 28 ± 1.249 | 27.56 ± 1.613 | 29.26 ± 3.447 | 31.89 ± 4.646 | 25.57 ± 5.223 |

### Global Metabolomic Profiling

Metabolites were extracted and analyzed by LC-MS analysis as previously described with slight modifications [21, 22]. SF samples were thawed on ice and centrifuged at 4°C at 500xg for 5 minutes to eliminate cells and debris. The supernatant was resuspended in 50:50 water:acetonitrile at −20°C for 30 minutes. The sample was vortexed for 3 minutes and centrifuged at 16100xg for 5 minutes at 4°C. The supernatant was completely evaporated in a vacuum concentrator for ∼2 hours, and the dried pellet was resuspended in 500 µL of acetone to precipitate proteins at 4°C for 30 minutes. The sample was then centrifuged at 16100xg for 5 minutes. The supernatant was completely evaporated by speedvac, and the pellet was resuspended in mass spectrometry grade 50:50 water:acetonitrile. Metabolite extracts were analyzed in positive mode using an Agilent 1290 UPLC system connected to an Agilent 6538 Q-TOF mass spectrometer (Agilent Santa Clara, CA). Metabolites were chromatographically separated on a Cogent Diamond Hydride HILIC 150×2.1 mm column (MicroSolv, Eatontown, NJ) using an optimized normal phase gradient elution method, and spectra were processed as previously described [18].

### Statistical Methods and Analysis

Global metabolomic profiling generates a large multivariate dataset of thousands of mass-to-charge ratios (m/z) and their corresponding peak intensities [17]. The dataset was reduced by removing metabolite features (m/z values) with median intensity values of zero across all experimental groups. All data analysis steps were completed using MetaboAnalyst unless otherwise noted [23]. Data were log transformed to correct for non-normal distributions and standardized (mean centered divided by standard deviation). Standardized data were used for all analyses unless indicated otherwise.

All statistical tests used an *a priori* significance level of 0.05, and false discovery rate (FDR) corrections were applied when performing multiple comparisons per metabolite between groups. The Kolomogorov-Smirnov test (KS-test) was used in MATLAB (MathWorks, Inc. Natick, MA) to compare cumulative median metabolite distributions between cohorts. This nonparametric test does not require assumptions about the underlying distributions and therefore is useful for metabolomics datasets that typically contain non-normal distributions. Specific differences between multiple groups were determined using analysis of variance (ANOVA) F-tests. Two-tailed Student’s t-tests examined specific differences between two groups only. Differentially regulated metabolites between two groups were visualized by volcano plot to assess both significance and magnitude of change simultaneously. Metabolite features with a p-value (FDR corrected) less than 0.05 and greater than twofold change were considered both statistically significant and biologically important in these analyses.

Multivariate methods assessed variations in the metabolomic datasets. Unsupervised hierarchical clustering analysis (HCA) based on Euclidean distance and average linkage separated samples into groups of similar abundance patterns [24]. HCA assessed subgroups of donors exhibiting distinct OA phenotypes. HCA is visualized using heatmaps, known as a clustergrams, to analyze the overall metabolomic profiles. Clustergrams reveal both clusters of co-regulated metabolite features and the relative similarity between experimental groups [24]. Principal component analysis (PCA) is another unsupervised method used to analyze metabolomics data. PCA orthogonally transforms a set of observations into principal components that each represent a fraction of the overall variance within the dataset. Partial least squares-discriminant analysis (PLS-DA) is a supervised classification method that reveals the underlying source of distinction between known groups. PLS-DA scores each variable in each component indicating how important that variable was in contributing to the separation.

Metabolite features (m/z values) were matched to known metabolite identities and mapped to relevant pathways using the metabolite library and pathway enrichment tool, *mummichog* [25]. *Mummichog* predicts a network of functional activity based on the projection of detected metabolite features onto local pathways. Pathway libraries MFN and Biocyc were used for compound identification and pathway enrichment (mass tolerance: 0.1 ppm; positive mode). Pathways reported were significant by pathway overrepresentation analysis with an FDR-adjusted p-value less than 0.05.

To determine if cohorts or phenotypes were associated with any confounding variables, Student’s t-tests, logistic regression, and post hoc Chi Squared tests were employed to assess differences between groups based on the available clinical data including age, sex, and BMI were assessed between both groups and phenotypes.

## Results

### Differences in Global Metabolomes between Healthy Donors, Early and Late OA

A total of 9903 metabolite features were detected in SF from donors with grade 0-IV OA. This dataset was refined to 1362 detected features by removing features with a median intensity of zero. ANOVA identified 39 differentially expressed metabolite features between healthy, early OA, and late OA SF (FDR-corrected p<0.05).

We first explored whether the global metabolomes were distinct between healthy, early, and late OA cohorts. To examine differences between cohorts, three pairwise comparisons were made: healthy vs. early OA; healthy vs. late OA; and early vs. late OA. Between-group differences in global metabolomes were assessed using KS-tests, and this revealed significant differences between all pairwise comparisons (p_ks_<0.01; Fig. 1). Taken together, these results indicate that the global metabolomes are significantly different between healthy, early, and late OA.

**Figure 1.**
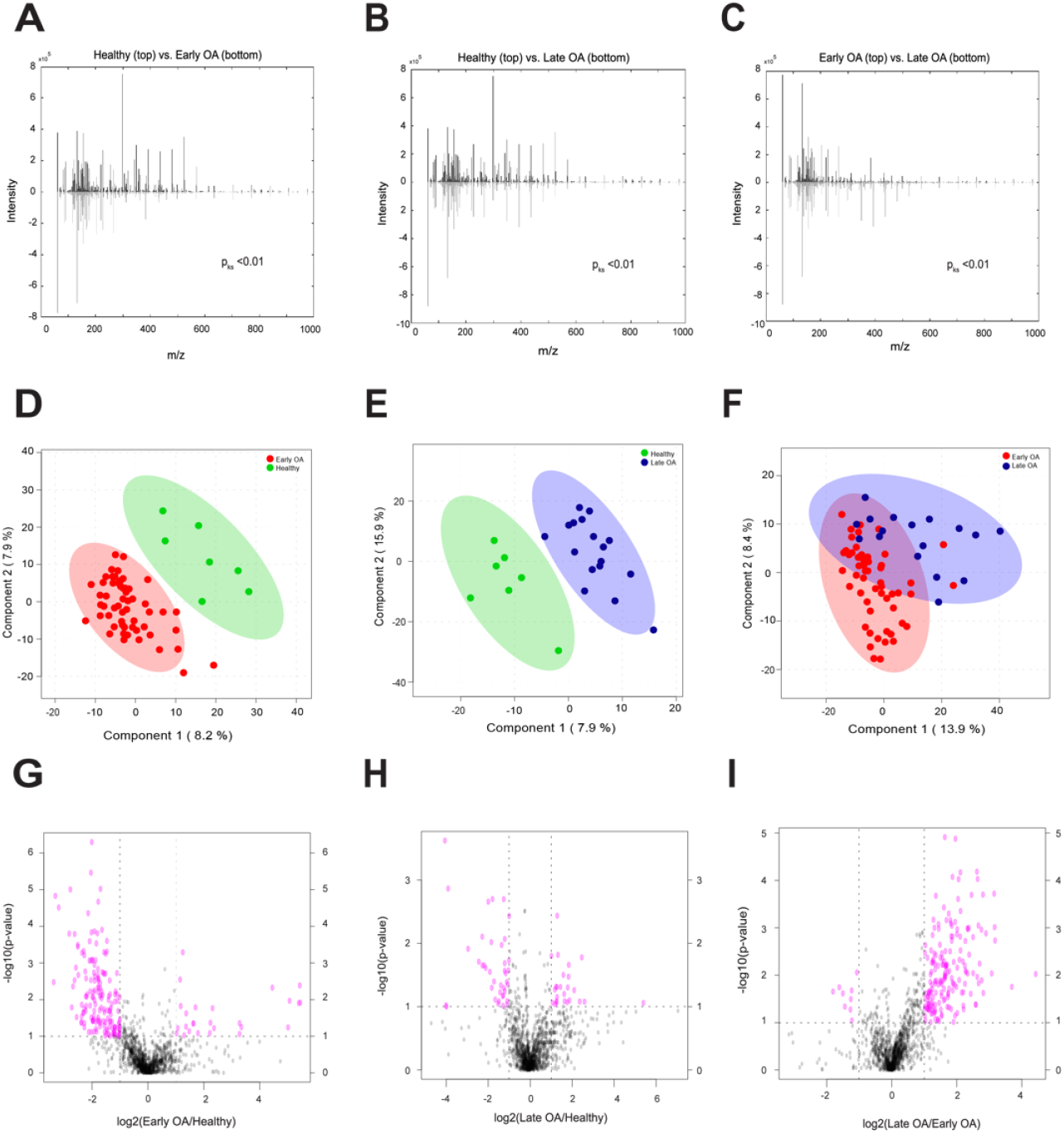
Global metabolomes are distinct between cohorts. **(A-C)** The cumulative distribution of metabolites between groups were distinct from one another. KS-tests comparing the median metabolite intensity distributions between groups revealed significantly (p_ks_<0.01) different metabolomic profiles. Mirrored metabolite distributions display differences between groups. **(D-F)** PLS-DA displayed differences in metabolomic profiles of between groups, revealing clear separation between healthy and OA donors and some separation between early and late OA donors. The first two components are plotted against one another with their contribution to the overall variance. 95% confidence ellipses illustrate class separation. **(G-I)** Volcano plot analysis between groups reveal metabolite features upregulated and downregulated by p-value and fold change analysis. Dashed lines indicate the p-value threshold of 0.05 (horizontal) and fold change threshold of 2 (vertical). The upper right and left quadrants contain significant (p<0.05) upregulated and downregulated features with a fold change greater than twofold. Metabolite features in the upper right and left quadrants were assessed for enriched pathways reported in Supplemental Table 2.

To visualize differences in metabolomic profiles and identify specific metabolite features with the greatest discriminative capabilities for separating cohorts, supervised PLS-DA was used. PLS-DA shows clear separation of healthy donors from disease donors, and minimal overlap between early and late OA donors (Fig. 1). By examining VIP scores, we found metabolite features that contribute the most to distinguishing between cohorts and are strong candidates for potential metabolite biomarkers (Supplemental Table 1).

Volcano plot analysis examined pairwise differences using both significance and fold changes (Fig. 1). 188 metabolite features were significantly different between healthy and early OA SF with 162 lower and 26 higher in early OA. 64 metabolite features were significantly different between healthy and late OA SF, with 39 lower and 25 higher in abundance. Within OA, 191 metabolite features were significantly different between early and late stage disease, with 9 lower and 182 higher in late stage disease. To infer metabolic activity, significantly different metabolite features were enriched using *mummichog’s* pathway analysis (Supplemental Table 2) presented below.

### Co-Regulated Metabolites Map to Differentially Regulated Metabolic Pathways with Disease

Early and late OA profiles were distinct from healthy SF (Fig. 2). Unsupervised HCA of healthy and diseased SF showed that the early and late OA profiles were more similar to one another than healthy SF (Supplemental Fig. 1). From the clustering, six groups of co-regulated metabolites were identified based on consistency of clustered distance and assessed for enriched pathways associated with stage of OA. (Supplemental Table 3).

**Figure 2.**
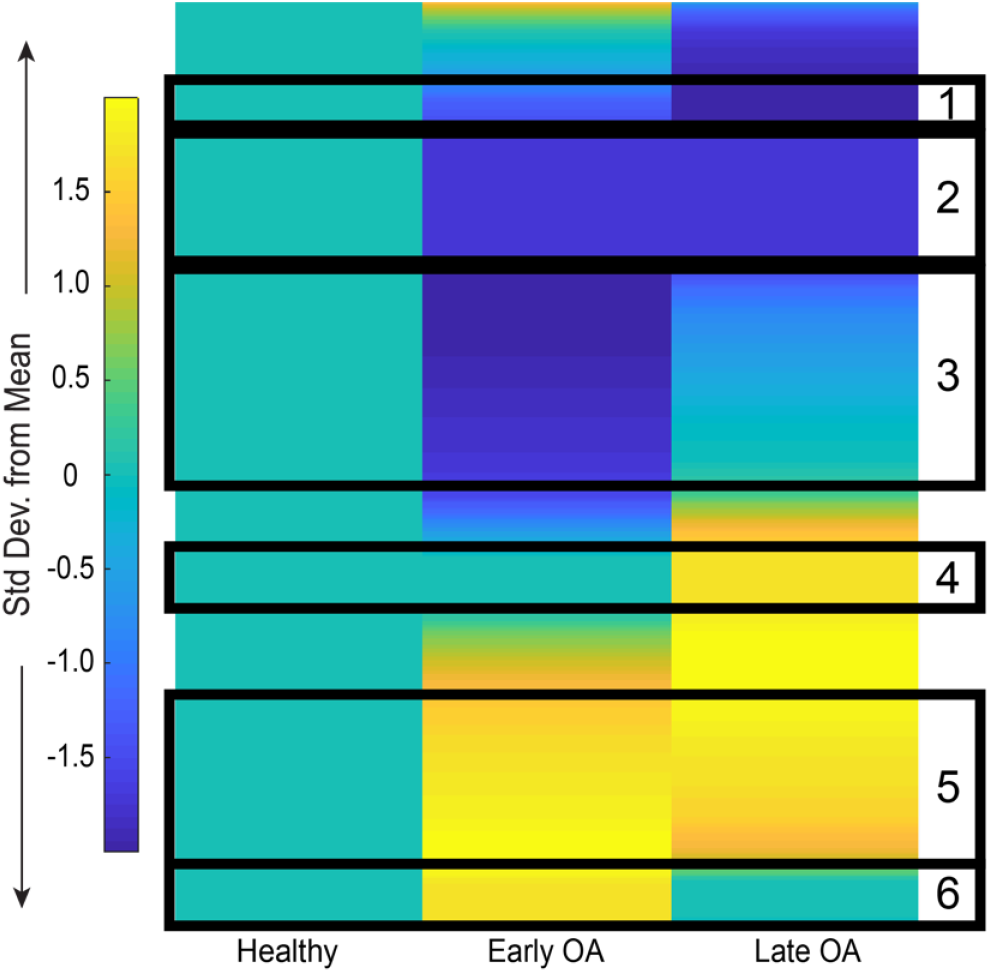
Metabolic changes in SF during early and late stage OA. Clustergram of median global metabolomic profiles of early and late OA SF normalized to healthy SF display patterns of metabolite expression with disease. Arbitrarily selected clusters of co-regulated metabolite features are boxed in black and enriched for relevant pathways in Supplemental Table 3.

Cluster 1 contained 38 metabolite features that decreased throughout disease progression. These mapped to 14 of the previously identified enriched pathways (Supplemental Table 2) including amino acid metabolism (glycine, serine, alanine, threonine, lysine, arginine, and proline), the urea cycle, phosphatidylinositol phosphate metabolism, the carnitine shuttle, vitamin metabolism (B5 and C), and porphyrin metabolism (Supplemental Table 3).

Cluster 2 contained 135 metabolite features that decreased in OA compared to healthy SF. These metabolite features mapped to 20 enriched pathways including vitamin metabolism (E, C, B3, and B6), phosphatidylinositol phosphate metabolism, glutathione metabolism, leukotriene metabolism, butanoate metabolism, amino acid metabolism (similar to cluster 1 with the addition of tryptophan and histidine metabolism), and the carnitine shuttle (Supplemental Table 3).

Cluster 3 contained 188 metabolite features lowest in early OA compared to healthy and late OA. These mapped to 14 enriched pathways including porphyrin metabolism, galactose metabolism, fructose and mannose metabolism, vitamin metabolism (B5, B3, E), methionine and cysteine metabolism, N-glycan degradation, glycerophospholipid metabolism, and leukotriene metabolism (Supplemental Table 3).

Clusters 4-6 contained metabolism features higher in abundance in OA cohorts. Cluster 4 contained 64 metabolite features highest in late OA. These metabolite features mapped to 8 enriched pathways including keratan sulfate degradation, N-glycan degradation, fructose and mannose metabolism, leukotriene metabolism, and butanoate metabolism (Supplemental Table 3).

Cluster 5 contained 177 metabolite features with the greatest abundance in early and late OA SF. These mapped to 36 enriched pathways including amino acid metabolism (histidine, glycine, serine, alanine, threonine, tyrosine, glutamate, aspartate, valine, leucine, isoleucine, aspartate, asparagine, lysine, and tryptophan) urea cycle, keratan sulfate degradation, fatty acid metabolism, glycerophospholipid and glycosphingolipid metabolism, the TCA cycle, N-glycan metabolism, glutathione metabolism, tryptophan metabolism, and vitamin C metabolism (Supplemental Table 3).

Cluster 6 contained 60 metabolite features highest in abundance in early OA. These mapped to 33 enriched pathways included glycolysis and gluconeogenesis, the pentose phosphate pathway, sialic acid metabolism, N-glycan degradation, keratan sulfate degradation, tryptophan metabolism, glutathione metabolism, and vitamin B3 metabolism (Supplemental Table 3).

### Unsupervised Clustering Suggests OA Phenotypes within Early and Late OA

To examine OA phenotypes, early and late OA were further analyzed by unsupervised HCA. In early OA, this revealed two clusters of donors, E1 and E2, containing 33 and 22 donors, respectively (Fig. 3A). There were 379 metabolite features differentially expressed between phenotypes E1 and E2 (FDR-corrected p<0.05). HCA of late OA also showed two distinct clusters of donors, L1 and L2, that may be representative of late OA phenotypes (Fig. 4A). 11 donors clustered in phenotype L1, and 6 donors clustered in phenotype L2. There were 187 differentially expressed metabolite features between phenotypes L1 and L2 (FDR-corrected p<0.05).

**Figure 3.**
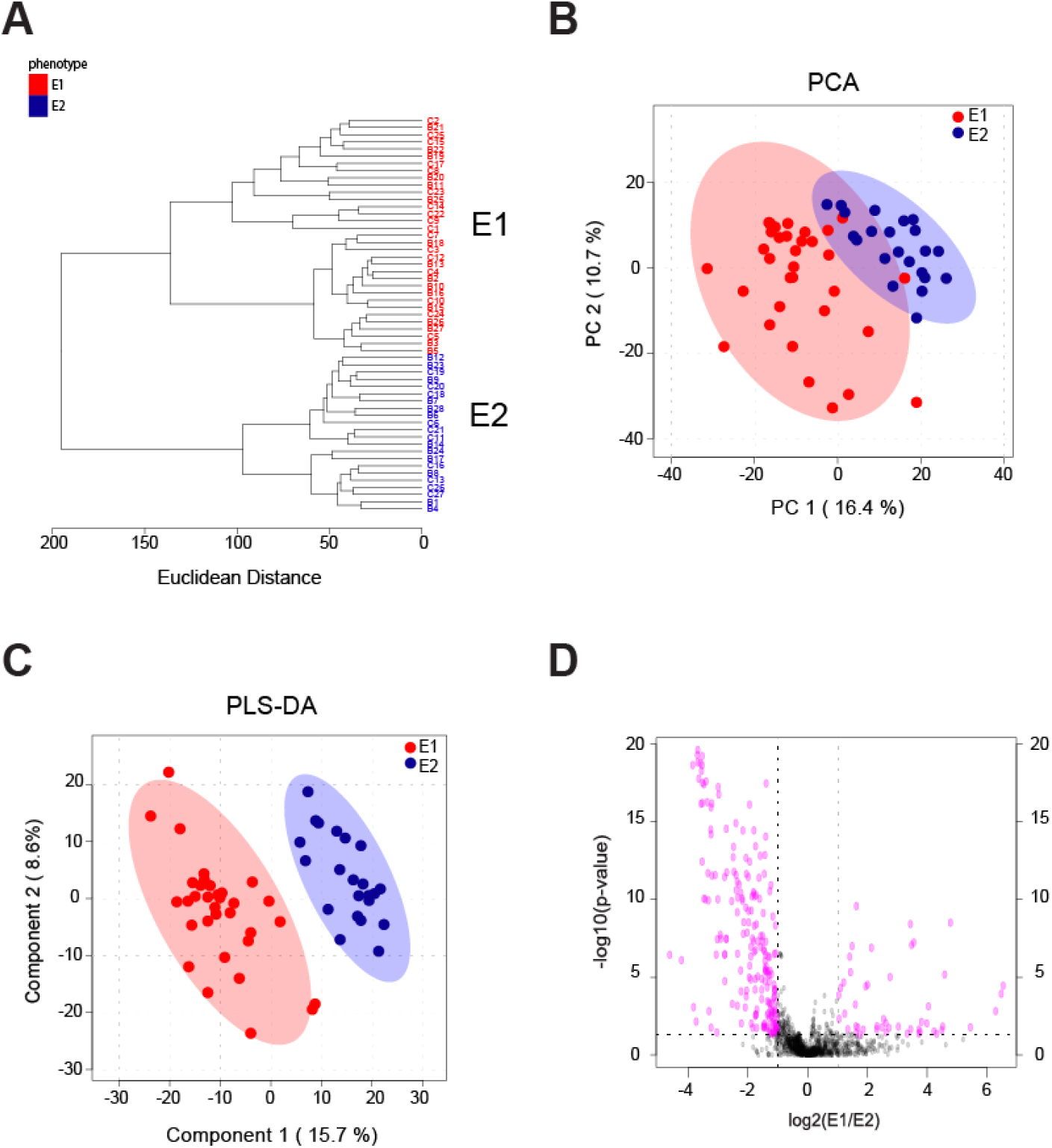
Phenotypes in early OA synovial fluid. **(A)** Unsupervised HCA of all early OA donors. Two clusters of donors were identified and labeled as phenotype E1 (red) and phenotype E2 (blue). E1 contained 33 donors and E2 contained 22. Line length represents Euclidean distances between donors and clusters. **(B)** Unsupervised PCA of all early OA donors reveals separation of early OA phenotypes. The first two components are associated with 27.1% of the variation between phenotypes. (E1=red; E2=blue). **(C)** Supervised PLS-DA further illustrated the separation between phenotypes (E1=red; E2=blue) with PC1 and PC2 accounting for 24.3% of the variance. **(D)** Volcano plot visualization of differentially regulated metabolite features by Student’s t-test significance and fold change analysis (E1:E2). The p-value threshold is represented by the horizontal dashed line (FDR-corrected p<0.05), and the vertical lines represent the fold change threshold (greater than twofold change). Metabolite features in the upper right and left quadrants (p<0.05 and greater than twofold change) were enriched for relevant pathways reported in Table 3, with the full list of perturbed pathways in Supplemental Table 2.

**Figure 4.**
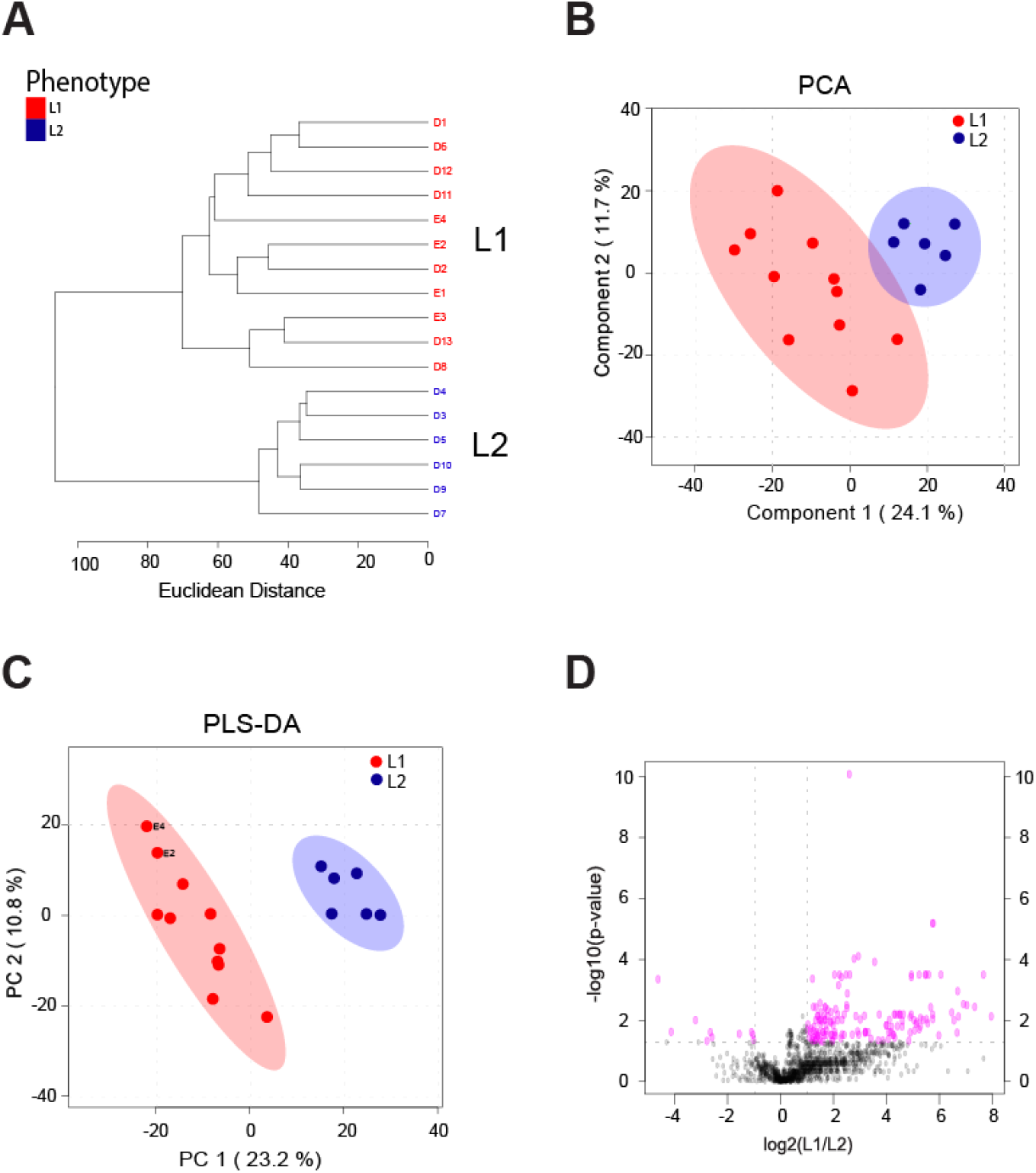
Phenotypes in late OA synovial fluid. **(A)** Unsupervised HCA of all late OA donors. Two clusters of donors were identified and labeled as phenotype L1 (red) and phenotype L2 (blue). L1 contained 11 donors, and L2 contained 6 donors. Line length represents Euclidean distances between donors and clusters. **(B)** Unsupervised PCA of all early OA donors reveals separation of early OA phenotypes. The first two PCs are associated with 35.8% of the variation between phenotypes. (L1=red; L2=blue). **(C)** Supervised PLS-DA further illustrated the separation between phenotypes (L1=red; L2=blue), with component 1 and component 2 accounting for 34% of the overall variance. **(D)** Volcano plot visualization of differentially regulated metabolite features by Student’s t-test significance and fold change analysis (L1:L2). The p-value threshold is represented by the horizontal dashed line (FDR-corrected p<0.05) and the vertical lines represent the fold change threshold (greater than twofold change). Metabolite features in the upper right and left quadrants were assessed for enriched pathways reported in Table 4.6.

PCA, an unsupervised method, was used to examine the separation between potential phenotypes. Plotting the PCA scores of early OA donors shows the separation between phenotypes, with PC1 and PC2 accounting for 27.1% of the overall variance (Fig. 3B). Separation of late OA donors into two distinct phenotypes is also supported by PCA, with PC1 and PC2 associated with 35.8% of the overall variance (Fig. 4B). PLS-DA, a supervised method, further supports distinct phenotypes within early and late OA as indicated by separation between E1 and E2 donors and L1 and L2 donors (Fig. 3C, 4C). Taken together, HCA, PCA, and PLS-DA support four distinct subgroups of donors in early and late stage disease that may be representative of metabolic OA phenotypes.

Distinct pathways were represented in the various phenotypes as determined by analyzing differentially expressed metabolites for enriched pathways. Volcano plot analysis found 254 metabolite features differentially expressed between the early OA phenotypes and 158 metabolite features differentially expressed between late OA phenotypes (Fig. 3D, 4D). Enrichment analysis was then employed to map differentially expressed metabolite features to pathways (Tables 2-3).

**Table 2.** Perturbed pathways in early OA phenotypes. Pathway enrichment of significant metabolite features upregulated and downregulated (p<0.05; greater than twofold change) with early OA phenotypes in Fig. 4.5F volcano plot analysis comparing phenotype E1 to phenotype E2 (E1:E2 fold change ratio). Significant metabolite features greater in abundance in the upper right quadrant of the volcano plot (higher in E1 compared to E2) in Fig. 3D were enriched to reveal corresponding upregulated pathways. Significant metabolite features reduced in abundance in the upper left quadrant of the volcano plot (lower in E1 compared to E2) in Fig. 3D were enriched to reveal corresponding downregulated pathways. Pathways are reported with the total metabolites in the pathway, the total detected metabolites in the pathway, and total significant (by volcano plot analysis) metabolites within that pathway. Only pathways with an FDR-corrected p-value less than 0.05 are reported. The full list of pathways identified in Fig. 3D volcano plot is reported in Supplemental Table 2.

| Downregulated in Early OA Phenotype E1 (Upregulated in Phenotype E2) |  |  |  |  |
| --- | --- | --- | --- | --- |
|  | Total | Detected | Significant | P-value |
| Phosphatidylinositol phosphate metabolism | 59 | 21 | 10 | 0.00031787 |
| Chondroitin sulfate degradation | 37 | 6 | 4 | 0.00046556 |
| Heparan sulfate degradation | 34 | 6 | 4 | 0.00046556 |
| Glycosphingolipid biosynthesis - ganglioseries | 62 | 7 | 4 | 0.00058063 |
| Galactose metabolism | 41 | 29 | 9 | 0.00066311 |
| Hexose phosphorylation | 20 | 16 | 6 | 0.0006945 |
| Glycosphingolipid biosynthesis - globoseries | 16 | 4 | 3 | 0.00070358 |
| N-Glycan Degradation | 16 | 5 | 3 | 0.0010502 |
| N-Glycan biosynthesis | 48 | 11 | 4 | 0.0018166 |
| Hyaluronan Metabolism | 8 | 2 | 2 | 0.001875 |
| Fructose and mannose metabolism | 33 | 21 | 6 | 0.001927 |
| Urea cycle/amino group metabolism | 85 | 39 | 9 | 0.0031932 |
| Keratan sulfate degradation | 68 | 4 | 2 | 0.0068439 |
| Sialic acid metabolism | 107 | 21 | 5 | 0.0068821 |
| Glycosphingolipid metabolism | 67 | 16 | 4 | 0.0082138 |
| Vitamin B9 (folate) metabolism | 33 | 11 | 3 | 0.010679 |
| Starch and Sucrose Metabolism | 33 | 11 | 3 | 0.010679 |
| Alanine and Aspartate Metabolism | 30 | 17 | 4 | 0.010787 |
| Tryptophan metabolism | 94 | 39 | 7 | 0.024073 |
| Selenoamino acid metabolism | 35 | 14 | 3 | 0.025485 |
| Amino sugars metabolism | 69 | 21 | 4 | 0.027862 |
| Xenobiotics metabolism | 110 | 34 | 6 | 0.029208 |
| Histidine metabolism | 33 | 15 | 3 | 0.032525 |
| Porphyrin metabolism | 43 | 15 | 3 | 0.032525 |
| Ascorbate (Vitamin C) and Aldarate Metabolism | 29 | 9 | 2 | 0.046412 |
| Upregulated in Early OA Phenotype E1 (Downregulated in Phenotype E2) |  |  |  |  |
|  | Total | Detected | Significant | P-value |
| Butanoate metabolism | 34 | 19 | 3 | 0.0012595 |
| Purine metabolism | 80 | 38 | 4 | 0.0022534 |
| Glutamate metabolism | 15 | 10 | 2 | 0.0027626 |
| Methionine and cysteine metabolism | 94 | 30 | 3 | 0.0051311 |
| Leukotriene metabolism | 92 | 16 | 2 | 0.0078615 |
| Lysine metabolism | 52 | 19 | 2 | 0.011907 |
| Tryptophan metabolism | 94 | 39 | 3 | 0.012978 |
| Valine, leucine and isoleucine degradation | 65 | 20 | 2 | 0.013498 |
| Amino sugars metabolism | 69 | 21 | 2 | 0.015211 |
| Bile acid biosynthesis | 82 | 27 | 2 | 0.028037 |
| Arginine and Proline Metabolism | 45 | 28 | 2 | 0.03058 |
| Galactose metabolism | 41 | 29 | 2 | 0.033231 |
| Pyrimidine metabolism | 70 | 32 | 2 | 0.041794 |
| Drug metabolism - cytochrome P450 | 53 | 34 | 2 | 0.04797 |

**Table 3.** Perturbed pathways in late OA phenotypes. Pathway enrichment of significant metabolite features upregulated and downregulated p<0.05; greater than twofold change) with late OA phenotypes in Fig. 4D volcano plot analysis comparing phenotype L1 to phenotype L2 (L1:L2 fold change ratio). Significant metabolite features greater in abundance in the upper right quadrant of the volcano plot (higher in L1 compared to L2) in Fig. 4D were enriched to reveal corresponding upregulated pathways. Significant metabolite features reduced in abundance in the upper left quadrant of the volcano plot (lower in L1 compared to L2) in Fig. 4D were enriched to reveal corresponding downregulated pathways. Pathways are reported with the total metabolites in the pathway, the total detected metabolites within the pathway, and total significant (by volcano plot analysis) metabolites within that pathway. Only pathways with an FDR-corrected p-value less than 0.05 are reported. The full list of pathways identified in Fig. 4D volcano plot is reported in Supplemental Table 2.

| Downregulated in Late OA Phenotype L1 (Upregulated in Phenotype L2) |  |  |  |  |
| --- | --- | --- | --- | --- |
|  | Total | Detected | Significant | P-value |
| Keratan sulfate degradation | 68 | 4 | 2 | 0.019848 |
| N-Glycan Degradation | 16 | 5 | 2 | 0.020384 |
| Sialic acid metabolism | 107 | 21 | 2 | 0.02995 |
| Galactose metabolism | 41 | 29 | 2 | 0.035387 |
| Upregulated in Late OA Phenotype L1 (Downregulated in Phenotype L2) |  |  |  |  |
|  | Total | Detected | Significant | P-value |
| Porphyrin metabolism | 43 | 15 | 9 | 0.00013083 |
| Alanine and Aspartate Metabolism | 30 | 17 | 7 | 0.00014903 |
| Vitamin B9 (folate) metabolism | 33 | 11 | 5 | 0.00018086 |
| Vitamin B3 (nicotinate and nicotinamide) metabolism | 28 | 12 | 5 | 0.00020224 |
| Urea cycle/amino group metabolism | 85 | 39 | 10 | 0.00021275 |
| Glutamate metabolism | 15 | 10 | 4 | 0.0003258 |
| Leukotriene metabolism | 92 | 16 | 5 | 0.00037052 |
| Arginine and Proline Metabolism | 45 | 28 | 7 | 0.00039868 |
| Methionine and cysteine metabolism | 94 | 30 | 7 | 0.00053156 |
| Histidine metabolism | 33 | 15 | 4 | 0.001096 |
| Glutathione Metabolism | 19 | 10 | 3 | 0.0016661 |
| Nitrogen metabolism | 6 | 4 | 2 | 0.0020684 |
| Beta-Alanine metabolism | 20 | 11 | 3 | 0.0023048 |
| Butanoate metabolism | 34 | 19 | 4 | 0.003009 |
| Lysine metabolism | 52 | 19 | 4 | 0.003009 |
| Aspartate and asparagine metabolism | 114 | 62 | 10 | 0.0030692 |
| Valine, leucine and isoleucine degradation | 65 | 20 | 4 | 0.0038276 |
| Phosphatidylinositol phosphate metabolism | 59 | 21 | 4 | 0.0048344 |
| Amino sugars metabolism | 69 | 21 | 4 | 0.0048344 |
| Glycine, serine, alanine and threonine metabolism | 88 | 36 | 6 | 0.0049222 |
| Hexose phosphorylation | 20 | 16 | 3 | 0.0094927 |
| Squalene and cholesterol biosynthesis | 55 | 10 | 2 | 0.020059 |
| Purine metabolism | 80 | 38 | 5 | 0.024465 |
| Biopterin metabolism | 22 | 11 | 2 | 0.025759 |
| Arachidonic acid metabolism | 95 | 22 | 3 | 0.032157 |
| Vitamin E metabolism | 54 | 12 | 2 | 0.032261 |
| Pyrimidine metabolism | 70 | 32 | 4 | 0.035863 |
| Tyrosine metabolism | 160 | 59 | 7 | 0.039574 |
| Xenobiotics metabolism | 110 | 34 | 4 | 0.046425 |
| Selenoamino acid metabolism | 35 | 14 | 2 | 0.047506 |

A subgroup of donors at each stage of OA (E2 and L2) exhibited evidence of glycosaminoglycan degradation and structural deterioration. E2 was associated with 25 significantly enriched pathways, including glycosaminoglycan degradation, sialic acid and N-glycan metabolism, tryptophan metabolism, and ascorbate metabolism (Table 2). L2 was associated with 4 significantly enriched pathways including keratan sulfate and N-glycan degradation, sialic acid metabolism, and galactose metabolism (Table 3).

The remaining OA phenotypes, E1 and L1, were associated with increased inflammation. Phenotype E1 was associated 14 significantly enriched pathways including metabolism of butanoate and leukotrienes—both of which play a role in inflammation (Table 2). L1 was associated with 30 significantly enriched pathways including arachidonic acid metabolism and leukotriene metabolism (Table 3). Phenotype L1 was also associated with glutathione metabolism, which may be suggestive of altered levels of oxidative stress (Table 3).

#### Confounding variables

We evaluated if differences in metabolomic profiles between healthy, early, and late OA were associated with age, sex, or BMI as possible covariates (Table 1). The ages and BMI of the healthy, early, and late OA cohorts were calculated and analyzed by Student’s t-test. Male:female ratios were analyzed by logistic regression and chi-squared tests. There were significant differences in ages between healthy, early, and late OA comparisons with early OA younger than late OA (p<0.05). However, there was little to no evidence of differences in BMI or male:female ratios (p>0.05). Therefore, any differences noted between cohorts besides being due to the presence or absence of OA may be associated with aging.

## Discussion

To our knowledge, this is the first study to use LC-MS-based global metabolomic profiling of human SF to study OA phenotypes. A single prior study used a targeted approach based on 186 metabolites for this same goal and found that acylcarnitine and free carnitine levels were significantly different between subgroups [9]. In contrast, the global approach used here removes bias by not excluding metabolites *a priori*. By focusing on all detected metabolites, this study produced a network of pathways perturbed with OA. These data provide greater understanding of disease pathogenesis, therapeutic targets, and insight for biomarker discovery.

1362 metabolite features were detected in human SF by LC-MS analysis, and global metabolomic profiles were generated for healthy, early OA, and late OA SF. OA was associated with altered extracellular matrix component metabolism (glucosamine and galactosamine biosynthesis, ascorbate metabolism, keratin sulfate metabolism, and N-glycan metabolism), amino acid metabolism, fatty acid and lipid metabolism (glycosphingolipid and glycerophospholipid metabolism, the carnitine shuttle), inflammation (leukotriene metabolism), central energy metabolism (glycolysis and gluconeogenesis, the TCA cycle), oxidative stress (vitamin E, glutathione metabolism), and vitamin metabolism (C, E, B1, B3, B6, and B9).

### Structural Deterioration

Diseased SF exhibited greater evidence of tissue damage compared to healthy SF. Keratan sulfate degradation, N-glycan degradation, sialic acid metabolism, and ascorbate metabolism were altered with OA. Keratan sulfate, chondroitin sulfate, and heparin sulfate are glycosaminoglycans (GAGs) that function as building blocks of articular cartilage. Their presence in the SF typically indicates increased cartilage turnover [26]. In OA, the articular cartilage is degraded reducing GAG content [27, 28]. These data are consistent with both synthesis and degradation of GAGs in the SF of both early and late stage donors. OA cartilage also exhibits collagen damage [29]. We identified hydroxyproline as a metabolite with the greatest ability in distinguishing early from late OA. Sialic acids and N-glycans are also important components of lubricin, a mucinous glycoprotein that lines the cartilage surfaces and acts as a lubricant [30]. These pathways were perturbed in diseased SF suggesting that the SF function in lubrication is compromised.

### Vitamin Metabolism and Oxidative Stress

The physiological significance of vitamins E, B5, and C may relate to their roles as antioxidants to counteract the increased oxidative stress in the joint during OA [31]. Additional results from diseased SF suggest oxidative stress included glutathione metabolism. Furthermore, vitamin B3 is also a required cofactor for the production of nitric oxide (NO) by NO synthase. NO has been shown to have both catabolic and protective effects in OA by modulating a variety of inflammatory and anti-inflammatory mediators [32]. Thus, altered vitamin B3 metabolism may drive NO-related changes during OA pathogenesis. The altered antioxidant metabolism exhibited in OA SF in this study further supports a role for oxidative stress in the development of OA [33].

### Phenotypes of OA in Synovial Fluid from Early and Late Stage Disease

OA is a heterogeneous disease with varying presentation. Because of this, we investigated if distinct metabolic phenotypes existed within OA SF (*i.e.* early vs. late or within each). We identified two distinct phenotypes in early OA—E1 and E2 and two in late OA—L1 and L2.

Both inflammation and structural degradation are involved in OA. In early OA, a subset of donors (E1) was associated with greater inflammation, while the remaining donors (E2) exhibited evidence of greater structural deterioration. Similarly, in late OA, phenotype L1 was associated with inflammation and oxidative stress while L2 was associated with structural deterioration products. These data suggest that inflammation and degradation may not be as closely correlated as expected. Furthermore, because of the close relationship between inflammation and pain [34, 35], the inflammatory phenotypes E1 and L1 may be associated with increased pain.

As in late OA phenotype L1, oxidative stress and inflammation have been extensively studied for their role in OA pathogenesis, yet both contribute to OA by promoting cartilage degradation [36]. Despite this, phenotype L1 exhibited reduced structural deterioration products in the SF compared to L2. This suggests a structural damage phenotype at both early and late stage disease, an inflammatory phenotype in early OA, and an inflammatory and oxidative stress phenotype at late stage disease. Overall, these findings further support the heterogeneous nature of OA and suggest stage-dependent phenotypes that may drive differences in symptoms.

### Limitations

This study has limitations and also opens opportunities for future research. First, the sample size for this study was relatively small (n=75). Some cohorts, such as healthy (grade=0), consisted of only 6 samples, whereas early OA contained 55. With a small sample size, it is unlikely that all metabolic phenotypes were represented. Furthermore, this sample did not contain complete clinical information. Age and sex were provided for all donors, BMI was provided for most, but others lacked cause of death, prior medical history, and/or ethnicity. Importantly, age was identified as a potential confounder in this study. Age-matching within experimental cohorts would avoid potential confounding by age. Targeting specific inflammatory metabolites or degradation products may yield further insight into OA phenotypes, and expanded sample sizes may allow detection of OA biomarkers.

## Conclusions

This is the first study to generate global metabolomic profiles of early and late OA SF and identify OA phenotypes within early and late OA cohorts. The identified pathways in early and late OA provide insight into disease progression and provide several molecular pathways to further investigate as biomarkers of OA and as targets for drug discovery. Furthermore, the identification of specific OA phenotypes supports the heterogeneity of disease. Expansion of this study will identify candidate biomarkers of early and late OA in human SF and within OA phenotypes.

## Acknowledgements

We acknowledge Kathryn Howe, for her assistance with sample preparation, the NSF for funding (CMMI 1554708 to RKJ) and the Montana State University Proteomics, Metabolomics, and Mass Spectrometry Facility for assistance with sample analysis. This facility is supported in part by funding from the Murdock Charitable Trust and NIH P20GM103474 of the IDEA program and AG007996.

---

**Supplementary Figure 1.**
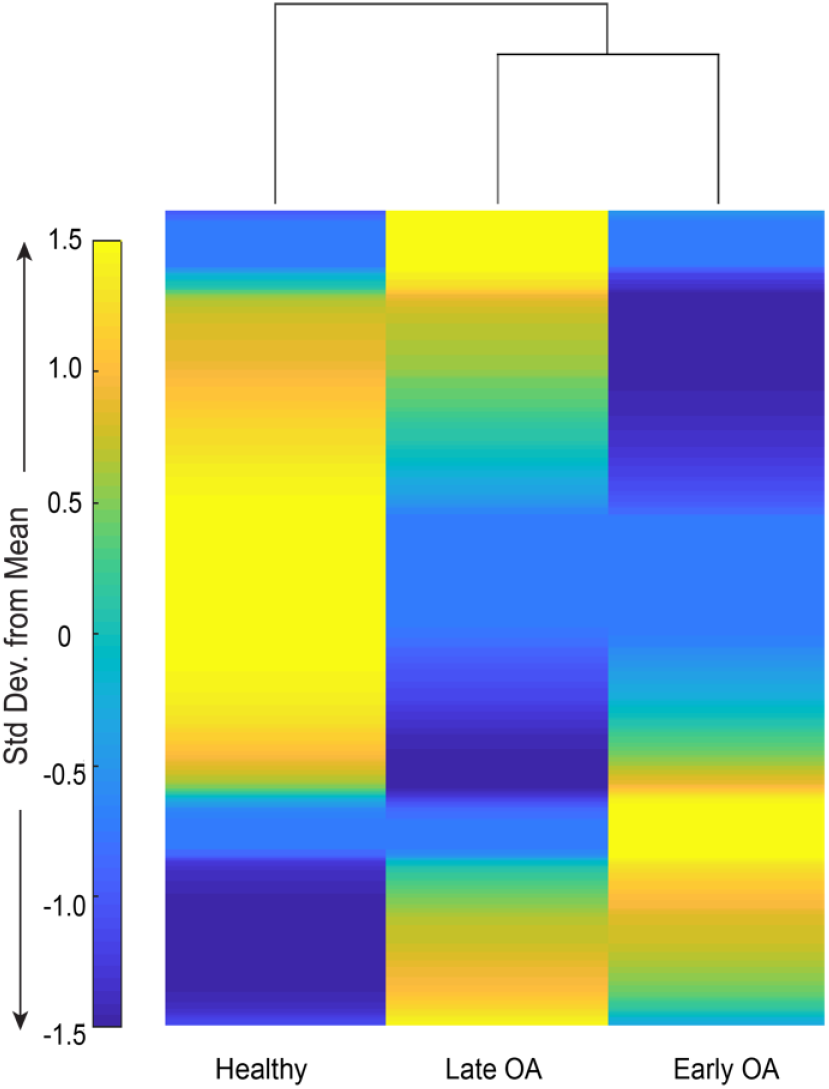
Distinct global metabolomic profiles of healthy, early OA, and late OA SF. Clustergram of median global metabolomic profiles of healthy, early OA, and late OA SF displays patterns of metabolite expression. HCA illustrates that early and late OA SF were more similar than healthy SF.

**Supplemental Table 1.** Discriminative metabolites identified by PLS-DA for classifying SF as healthy, early OA, late OA, phenotype E1, phenotype E2, phenotype L1, or phenotype L2. Full list of discriminative metabolite features with VIP scores for the top two components for each PLS-DA plot in Figure 1D-F, 3C, and 4C. Potential metabolite identities are reported as compound matches using *mummichog* and the Biocyc pathway library.

**Supplemental Table 2.** Distinct pathways are perturbed between groups. Full pathway enrichment of volcano plots in Fig. 1G-I, 3D, and 4.D. Pathways are reported for the pathway library, MFN.

**Supplemental Table 3.** Full list of all pathways identified for each cluster in Fig. 2 clustergram.

## References

1. Disease, G.B.D., I. Injury, and C. Prevalence, Global, regional, and national incidence, prevalence, and years lived with disability for 328 diseases and injuries for 195 countries, 1990-2016: a systematic analysis for the Global Burden of Disease Study 2016. Lancet, 2017. 390(10100): p. 1211–1259.

2. Hunter, D.J., et al., Biomarkers for osteoarthritis: current position and steps towards further validation. Best Pract Res Clin Rheumatol, 2014. 28(1): p. 61–71.

3. Loeser, R.F., et al., Osteoarthritis: a disease of the joint as an organ. Arthritis Rheum, 2012. 64(6): p. 1697–707.

4. Kulkarni, K., et al., Obesity and osteoarthritis. Maturitas, 2016. 89: p. 22–8.

5. Guilak, F., Biomechanical factors in osteoarthritis. Best Pract Res Clin Rheumatol, 2011. 25(6): p. 815–23.

6. Kramer, W.C., K.J. Hendricks, and J. Wang, Pathogenetic mechanisms of posttraumatic osteoarthritis: opportunities for early intervention. Int J Clin Exp Med, 2011. 4(4): p. 285–98.

7. Li, Y., et al., The age-related changes in cartilage and osteoarthritis. Biomed Res Int, 2013. 2013: p. 916530.

8. Deveza, L.A., et al., Knee osteoarthritis phenotypes and their relevance for outcomes: a systematic review. Osteoarthritis Cartilage, 2017. 25(12): p. 1926–1941.

9. Zhang, W., et al., Classification of osteoarthritis phenotypes by metabolomics analysis. BMJ Open, 2014. 4(11): p. e006286.

10. Karsdal, M.A., et al., OA phenotypes, rather than disease stage, drive structural progression--identification of structural progressors from 2 phase III randomized clinical studies with symptomatic knee OA. Osteoarthritis Cartilage, 2015. 23(4): p. 550–8.

11. Bartlett, S.J., et al., Identifying common trajectories of joint space narrowing over two years in knee osteoarthritis. Arthritis Care Res (Hoboken), 2011. 63(12): p. 1722–8.

12. Collins, J.E., et al., Trajectories and risk profiles of pain in persons with radiographic, symptomatic knee osteoarthritis: data from the osteoarthritis initiative. Osteoarthritis Cartilage, 2014. 22(5): p. 622–30.

13. Karsdal, M.A., et al., Disease-modifying treatments for osteoarthritis (DMOADs) of the knee and hip: lessons learned from failures and opportunities for the future. Osteoarthritis Cartilage, 2016. 24(12): p. 2013–2021.

14. Castaneda, S., et al., Osteoarthritis: a progressive disease with changing phenotypes. Rheumatology (Oxford), 2014. 53(1): p. 1–3.

15. Bierma-Zeinstra, S.M. and A.P. Verhagen, Osteoarthritis subpopulations and implications for clinical trial design. Arthritis Res Ther, 2011. 13(2): p. 213.

16. Patti, G.J., O. Yanes, and G. Siuzdak, Innovation: Metabolomics: the apogee of the omics trilogy. Nat Rev Mol Cell Biol, 2012. 13(4): p. 263–9.

17. Gowda, G.A. and D. Djukovic, Overview of mass spectrometry-based metabolomics: opportunities and challenges. Methods Mol Biol, 2014. 1198: p. 3–12.

18. Carlson, A.K., et al., Application of global metabolomic profiling of synovial fluid for osteoarthritis biomarkers. Biochem Biophys Res Commun, 2018. 499(2): p. 182–188.

19. Brandt, K.D., The role of analgesics in the management of osteoarthritis pain. Am J Ther, 2000. 7(2): p. 75–90.

20. Outerbridge, R.E., The etiology of chondromalacia patellae. J Bone Joint Surg Br, 1961. 43-B: p. 752–7.

21. Carlson, A.K., et al., Application of global metabolomic profiling of synovial fluid for osteoarthritis biomarkers. Biochem Biophys Res Commun, 2018.

22. Kim, S., et al., Global metabolite profiling of synovial fluid for the specific diagnosis of rheumatoid arthritis from other inflammatory arthritis. PLoS One, 2014. 9(6): p. e97501.

23. Xia, J. and D.S. Wishart, Using MetaboAnalyst 3.0 for Comprehensive Metabolomics Data Analysis. Curr Protoc Bioinformatics, 2016. 55: p. 14 10 1–14 10 91.

24. Bar-Joseph, Z., D.K. Gifford, and T.S. Jaakkola, Fast optimal leaf ordering for hierarchical clustering. Bioinformatics, 2001. 17 Suppl 1: p. S22–9.

25. Li, S., et al., Predicting network activity from high throughput metabolomics. PLoS Comput Biol, 2013. 9(7): p. e1003123.

26. Thonar, E.J., et al., Keratan sulfate in body fluids in joint disease. Acta Orthop Scand Suppl, 1995. 266: p. 103–6.

27. Elliott, R.J. and D.L. Gardner, Changes with age in the glycosaminoglycans of human articular cartilage. Ann Rheum Dis, 1979. 38(4): p. 371–7.

28. Hjertquist, S.O. and R. Lemperg, Identification and concentration of the glycosaminoglycans of human articular cartilage in relation to age and osteoarthritis. Calcif Tissue Res, 1972. 10(3): p. 223–37.

29. Hosseininia, S., L.R. Lindberg, and L.E. Dahlberg, Cartilage collagen damage in hip osteoarthritis similar to that seen in knee osteoarthritis; a case-control study of relationship between collagen, glycosaminoglycan and cartilage swelling. BMC Musculoskelet Disord, 2013. 14: p. 18.

30. Jay, G.D., et al., Prevention of cartilage degeneration and gait asymmetry by lubricin tribosupplementation in the rat following anterior cruciate ligament transection. Arthritis Rheum, 2012. 64(4): p. 1162–71.

31. Collins, J.A., et al., Oxidative Stress Promotes Peroxiredoxin Hyperoxidation and Attenuates Pro-survival Signaling in Aging Chondrocytes.J Biol Chem, 2016. 291(13): p. 6641–54.

32. Abramson, S.B., Osteoarthritis and nitric oxide. Osteoarthritis Cartilage, 2008. 16 Suppl 2: p. S15–20.

33. Loeser, R.F., et al., Aging and oxidative stress reduce the response of human articular chondrocytes to insulin-like growth factor 1 and osteogenic protein 1. Arthritis Rheumatol, 2014. 66(8): p. 2201–9.

34. Ballegaard, C., et al., Knee pain and inflammation in the infrapatellar fat pad estimated by conventional and dynamic contrast-enhanced magnetic resonance imaging in obese patients with osteoarthritis: a cross-sectional study. Osteoarthritis Cartilage, 2014. 22(7): p. 933–40.

35. Miotla Zarebska,J., et al., CCL2 and CCR2 regulate pain-related behaviour and early gene expression in post-traumatic murine osteoarthritis but contribute little to chondropathy. Osteoarthritis Cartilage, 2017. 25(3): p. 406–412.

36. Lepetsos, P. and A.G. Papavassiliou, ROS/oxidative stress signaling in osteoarthritis. Biochim Biophys Acta, 2016. 1862(4): p. 576–591.

